# RNA and proteins joined up at the origins of life: Persistence is the point

**DOI:** 10.64898/2026.07.09.737588

**Authors:** Mohamed Swailem, Ken A. Dill

## Abstract

What drove nucleic acids (NA) into functional associations with proteins (PR) at the origins of life? We reason from polymer physics and the Central Dogma that the fitness value of cooperating through a division of labor – NA for replication fidelity and PR for functional fitness – is much higher than for either polymer alone. Our model shows a Pareto Front, where NA and PR can bootstrap each other to achieve autocatalytic cooperativity towards biology.

---

To understand how the first cells arose from prebiotic chemistry requires making sense of what drove the proteins and RNA/DNA molecules toward cooperative functioning (1, 2). Today’s living systems depend on both types of sequence-defined polymers: nucleic acids (NA) (RDNA, our shorthand for RNA and DNA) and protein molecules (PR). We call this a 2-polymer world (2PW). Could a simpler, or earlier, form of life have been a one-polymer world (1PW)? For example the *RNA World Hypothesis* says that because RNA molecules can both carry information and catalyze some reactions, RNA alone might have initiated the first life (3–6). Alternatively, a proteins world could have produced diverse functionalities prior to heritability (7–14).

The puzzle is framed by the Eigen paradox (15, 16), which we paraphrase as: *Good RNA is needed to make proteins but good proteins are needed to make RNA*. So naively, it would seem that originating a 2PW would be “twice as improbable” as a 1PW. However, that logic neglects fitness, where some PR–NA association has greater advantage than either individual polymer type alone. Here, we model the emergence of the two-polymer association based on a physical cooperativity motivated by the central dogma. The model shows that the fitness advantage of separation of labor – proteins impart functional fitness while RDNA imparts heritability – is so high that the event of their joining together could define the origins of life. To set the stage for our modeling, we first consider how the nature of evolution constrains possible relationships between the two polymer types.

## Biological evolution is the context that defines fitness

Biological evolution is a process of stochastic optimization performed by molecules. Fig. 1 shows evolution’s process: (1) Given a wild-type population x, mutate a cell, (2) grow up the mutant into a population y, (3) compete x vs. y for their relative abilities to grow and reproduce, (4) the winner gets more resources going forward into the next cycle. The relative ability to win additional resources is the fitness. Below, we first describe how the different physics of RDNA and proteins serve well the two different functions – namely to explore and exploit – needed for such optimization. Then, we model how their joint fitness is greater than that of the individual polymers.

**Figure 1:**
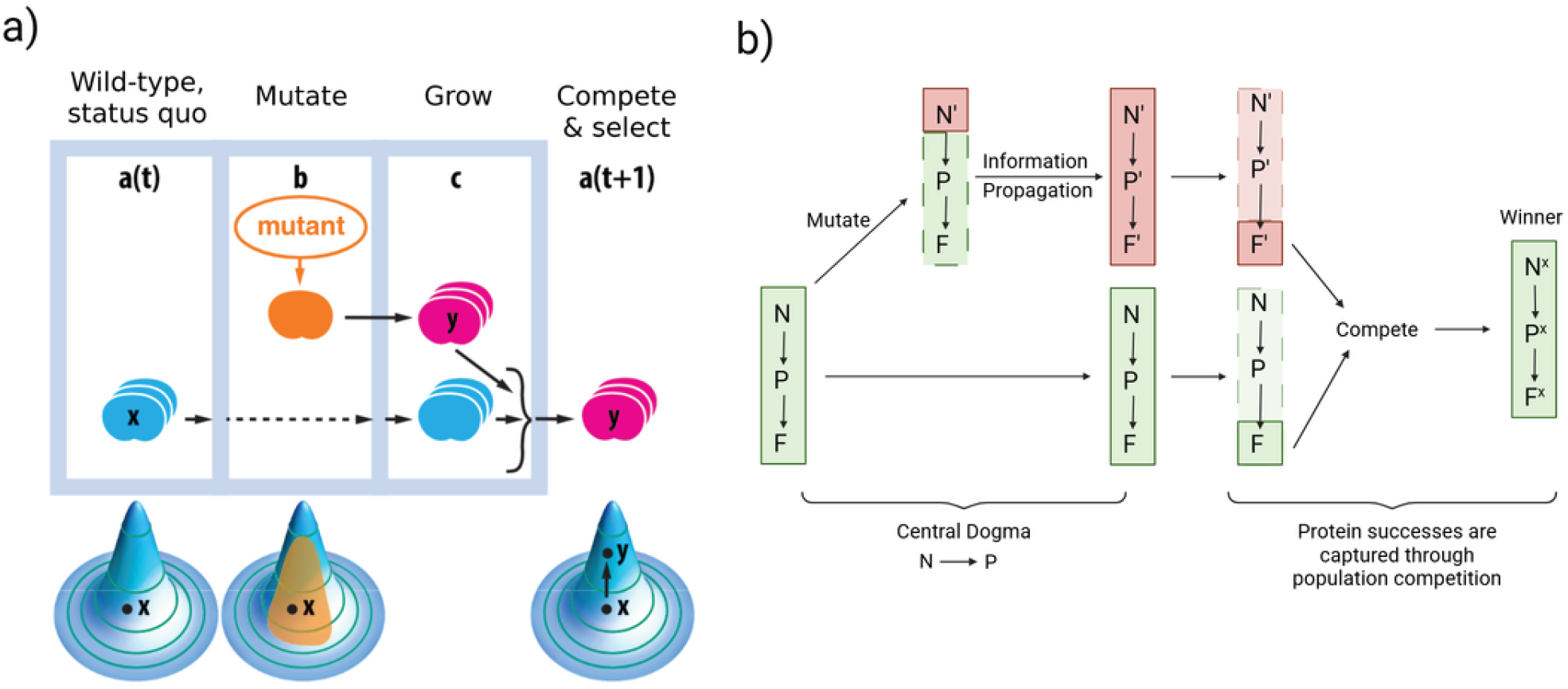
**a) Biological evolution as stochastic optimization:** Initially, wild-type cells *x* grow up a population. A mutant *y* grows up a population. The populations x and y compete. The winner (greater population) then fetches more resources. Fitness landscapes at the bottom show how the mutant is an experiment, where the winner decides the next step direction. **b) Evolution annotated in terms of the central dogma:** (Left, green) Wildtype starting point, NA encodes PR, which gives fitness F. (Top, red), Mutation in the blueprint NA, becoming NA’. (Red, green) both NA sequences grow up into populations. (Next) The populations compete for fitness. The winner, *F*^*x*^ now carries with it the winning sequence *NA*^*x*^ and protein *PR*^*x*^.

### The central dogma gives the relationship, *NA* → *PR*

The central dogma of molecular biology (CD) expresses a unidirectional flow of information, *NA → PR*, between the two polymer types; see Fig. 2. First elucidated by Francis Crick around 1970 (17), the central dogma is not simply that DNA makes RNA makes protein. Rather, Crick distinguished the informational flow from the mechanistic actions (18). The “informational” processes are of NAs mutating and translating into proteins inside a cell. The “action” processes are competition and selection outside the cell. The CD says that the informational flow is unidirectional; no protein information flows directly back into the NAs within the cell. The feedback loop is closed only through actions outside the cell, as the competition-winning cells increase their populations and their access to resources.

**Figure 2:**
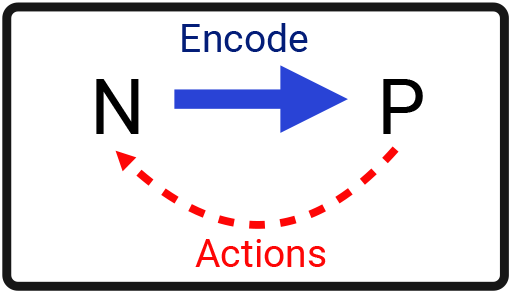
Central Dogma defines a directionality. Information flows in only one direction, namely from NA to PR, inside the cell (mutation and translation, blue arrow). Only through the external actions do the protein actions feedback into NAs (competition and selection, red arrow).

Fig. 1b gives more mechanistic detail. It shows *NA* as the blueprint and *PR* as the realization of the blueprint, resulting in fitness *F* of the wildtype. Information flows from *NA → PR*, and not the reverse. Mutations in the RDNA can alter proteins, but on the whole, the firewall of cellular machinery prevents protein errors to flow directly back.

To illuminate the CD mechanism, here is a metaphor. In an assembly line, the blueprint (nucleic acids) direct the making of product (proteins), resulting in some cost-benefit fitness F for the assembly line of the “factory” (the cell). Changes to the assembly line are performed in a particular way: First a random alteration in the blueprint from NA to NA^*′*^ is proposed; that directs an altered product from PR to PR^*′*^; and that, in turn, gives an altered fitness from F to F^*′*^. Each assembly line has its own blueprint attached to it at each step of the process. Then there is selection of whichever of the two assembly lines, call it *x*, has the better fitness, F^*x*^. That cycle of evolution/innovation repeats. Each step of each cycle carries “full documentation”, i.e. its own blueprint NA^*x*^ and its own PR^*x*^. Once the better assembly line is chosen, its blueprint is already carried along; see Fig. 1b. An alternative procedure would be to edit the blueprint after the winner is determined; that would allow for Lamarckian evolution see SI, but would violate the central dogma and is not consistent with Darwinian evolution.

The central dogma is a statement about today’s biology, not necessarily about how biology arose, or why, or if there are alternatives. But below, we argue that the CD relationship likely reflects important underlying physical constraints on how stochastic optimization can be implemented using molecules.

### Evolution’s stochastic optimization: a division of labor

The Darwinian evolution process may have emerged from something different and simpler, but ultimately the better stochastic optimization process will win out over worse ones. Examples of stochastic optimization computer algorithms are Monte Carlo, simulated annealing, and genetic algorithms. The Metropolis Monte Carlo (MMC) algorithm: make a random change to a state; compare it to the original; select the better of the two. There are two types of stochastic optimization processes: (i) when a fitness function and its gradient are available a priori and can be exploited by the optimization procedure or (ii) when the fitness uphill direction is not known, and must be discovered by trial and error. Evolution is of the latter type; fitnesses become known only through competitions and selection.

One of the deep principles of stochastic optimization is its separation into distinct actions, information-gathering and information-using components, called explore vs. exploit. Consider the MMC method. There are three components in taking a step: **(i) Propose:** A uniformly random “dice roll” chooses a possible step even-handedly drawn from a full space of alternative options. **(ii) Compete:** That potential step is then compared to the current state, the status quo value of the property of interest. **(iii) Select:** Accept the step or not based on a criterion of fitness or energy or cost-benefit function of some kind. These three components entail a division of labor that is essential to its success; namely of an ecumenical action (universal, even-handed, unbiased) versus a parochial action (specific, deciding, choosing, imparting bias).

The following conditions are necessary for a maximally innovative evolutionary process: **(1) The propose step must be ecumenical/unbiased**, showing no preference of one proposed option over any other. Every sequence in the whole space must be a possible option able to be considered without bias relative to every other sequence, otherwise the process would not be maximally innovative and exploratory. **(2) The select step must be parochial/biased**, meaning that it is evaluative, testing a specific fitness task, different for different sequences. The select step must strongly depend on the fitness landscape starting from the given point of the wild-type organism, while the propose step must be as completely independent of the landscape shape as is possible. We argue below that today’s biology is able to separate ecumenical from parochial actions by parsing different actions into the two different polymer types, NA and PR, because of their very different polymer physics.

### Evolution is stochastic optimization implemented in polymers

Biology’s sequence-defined polymers are suited for their different roles in this process; see Fig. 3.

**Figure 3:**
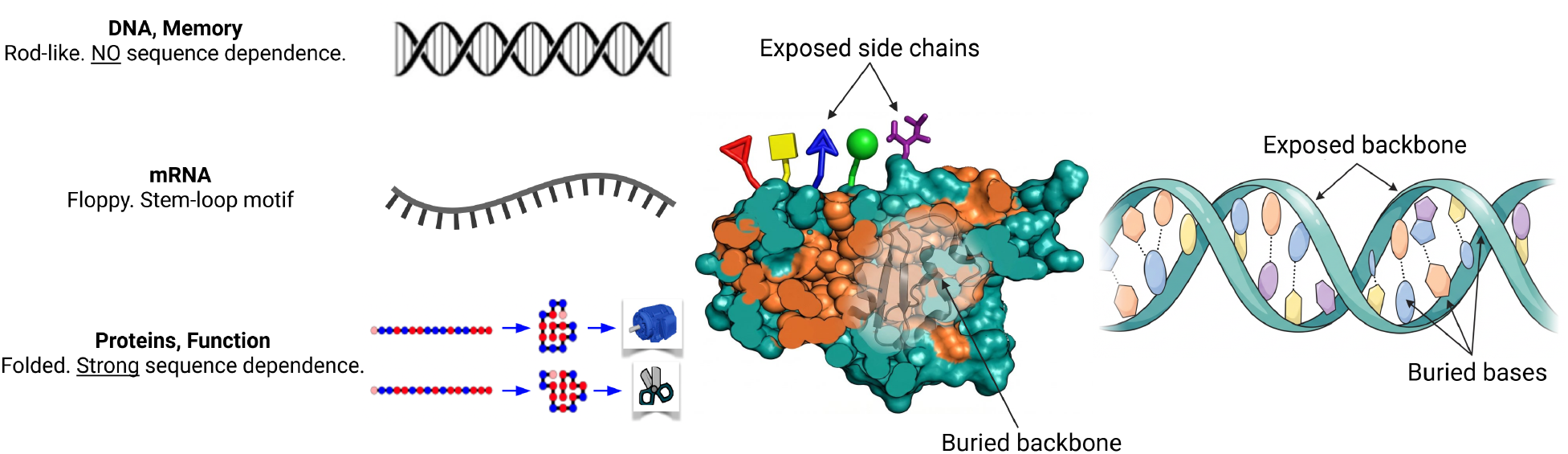
The polymer properties are quite different in DNA, RNA and proteins. DNA molecules are good at unbiased information storage, because the stiff double helical structure is agnostic to sequence and because their bases are buried inside, away from influence by external factors. RNAs, as mRNAs are mostly floppy, are well suited for carrying short stretches of sequence. Proteins are good at function because: they fold; folds are functional; they have 20 different chemical reaction moieties in the sidechains; and the sidechain gardens are exposed externally.

- **DNA is ecumenical, well-suited for read/ write/ memory/ bookkeeping**. DNA is an ideal blueprint material. As a form of matter, DNA is ecumenical; it can carry any sequence without significant preferential bias of one sequence over another. This is because DNA is stiff due to its double-helix structure, and because DNA bases are buried in the interiors of the double helix, relatively isolated from external environment effects. But correspondingly, DNA cannot replace protein folding and function, which are highly dependent on sequence.
- **Proteins are parochial, well-suited for encodable functionalities/ actions/ growth/ differential populations**. As a form of matter, amino acid sequences uniquely encode native structures and actions. Large fractions of protein sequence space appear capable of unique folding, including those in random libraries (11–13, 19). But correspondingly, proteins cannot substitute for RDNA. Proteins are largely unreadable and unwriteable; they would be poor as blueprints. Because most proteins are folded, trying to unfold and read them would be like trying to read a book that has its pages glued shut. And, because different proteins have different folded states, reading and writing in proteins would have to contend differently with every sequence.

What about amyloids or protein aggregates as information stores? For example, prions can induce copies (20, 21), or proteins involved in recognition through binding (22–24)? These are not substitutes for he information needed for evolution because they are not processes that can read or write individual sequences, nor are they applicable to arbitrary sequences.

- **RNA is neither**. Single-stranded RNAs are not ecumenical; their conformations depend on their sequences. They are also not parochial, because as a form of matter, most RNA molecules are floppy: they have 7 backbone degrees of freedom between monomers, vs 3 in proteins. RNAs typically have bumpy folding landscapes with many stable and metastable structures (25–27). RNAs are prone to kinetic trapping and rarely have a single folded structure. Rather, RNAs are often mRNAs, which are small floppy information carriers from the DNA repository to the ribosomal protein makers; see Fig. 3: (i) RNAs have mainly only 2 types of base pairs, while proteins have 20 different chemistries in their sidechains. (ii) RNA interactions are internal, in base pairing, not external, interacting with the outside world like proteins (Fig. 3). (iii) Because they tend to have multiple metastable states, a sequence doesn’t map to a single structure (28–30). (iv) RNA folding is dominated by secondary interactions favoring linear low-branching stem-loop motifs (31). The most common RNA functional molecules in most cells are ribosomal RNAs or tRNAs, both of which depend on protein partners.
- **Polysaccharides are not linearly encodable**. Sugar chains can carry information in their sequences of monomer types. But, they also have additional degrees of freedom – branching, linkage position and anomeric configuration (*α* or *β*). So, unlike RDNAs, polysaccharides have no template-based way of forming a linear information pipeline that can store → recover → copy. And unlike proteins, polysaccharides mostly don’t have unique folds and functions because of their hydrophilic character. So, sugar chains are neither ideal for read/write or folding/function actions.

Summarizing and simplifying: NA and PR play different roles, and each is well-suited for its niche. DNA is good at read/write/copy/memory, but not folding and function. Proteins are good at folding and function, but poor at read/write/information storage. These properties coincide with the need for a sequence-independent ecumenical mutate/propose step and a parochial compete/select step. Therefore, a compelling case can be made that life’s origins required a world of both polymers.

#### Origins was a transition from macro to micro catalysts

Life may have begun on a geographically localized macroscale catalyst that performed a particular reaction under given conditions – a volcanic vent, black smokers, a warm little pond, or a mineral or clay surface, for example (32–35). We call this macro fountainhead “a Founding Rock in Nebraska”. However, the transition to biology arguably required instead catalysis at the microscale – today mainly by protein molecules – that is molecular, mobile, editable for different catalytic actions and tailorable for different conditions (36). Below, we reason that the transition to such microscopic fountainheads required both RDNA and proteins, for both heritability and functional variability, perhaps in biomolecular condensates.

#### Origins was a transition of timescales, from individuals to lineages

Chemistry to biology was a transition of timescales. In chemistry, individual molecules decay, degrade, dilute or relax to equilibrium on molecular timescales. In contrast, evolution’s timescale is not lifetimes of individuals; it is the lifetimes of lineages of individuals and their progeny. Lineages are long-lived because of heritable propagation of short-lived individuals who come and go into or out of the lineage. Individuals are interchangeable. Lineages are about heritability. Oil droplets can divide, but they have no identifiable ancestry, hence no lineages. The transition from chemistry to biology was a transition from short molecule persistence times to long lineage persistence times.

### The transition from chemistry to biology requires cooperativity

Non-equilibrium systems, like living organisms, are often treated by birth-death dynamics, where a population *P* can be described by an equation *dP/dt* = *g*(*P*)*P − DP* wherein *g*(*P*) is the growth rate and *D* is the death rate. In a prebiotic version of this, there must be some process of molecules making more molecules; and decaying, breaking down, or diffusing away at a total rate *D*. The simplest starting point is a linear model, having a constant growth rate *g*(*P*) = *g*_1_ (a linear growth function), giving the following dynamics:

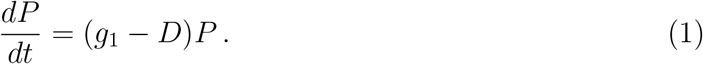

This equation describes the short-time transient after the discovery of an autocatalyst. Consider three regimes describing the fate of this autocatalyst. (1) Degradation wins and (*g*_1_ *− D*) *<* 0. For example, if the autocatalyst is thermodynamically unfavorable and the system is at thermodynamics equilibrium. In order for the autocatalyst’s growth to be sustained there needs to be some form of non-equilibrium drive leading to (2) Growth wins and (*g*_1_ *− D*) *>* 0 as long as the autocatalyst is in this favorable environment. (3) For the transition from chemistry to biology, this linear model is evidently not sufficient to explain a change in the sign of *dP/dt*. Taking the next lowest order in the short-time transient approximation is required to explain the transition from death-dominated to growth dominated. For more details on this invasion analysis see (8, 37). This leads to the following model:

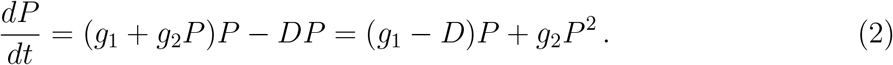

We refer to the *g*_2_ term as *cooperativity* (2) is not a mechanistic description; it is just a statement of what population dynamics would be required to transition from one regime (prebiotic, where death dominates) to another (postbiotic, where births dominate); see Fig. 4. In order to be a viable explanation of the transition to biology, a mechanism must do more than simple autocatalysis, more than just each dynamical agent making copies of itself. Along a reaction coordinate from chemistry to biology, there must have been some super-catalysis, some growing advantage of growth vs death terms. Below, we model how RDNA–protein association could originate a Central Dogma relationship by combining heritability or RDNA with function and fitness of proteins.

**Figure 4:**
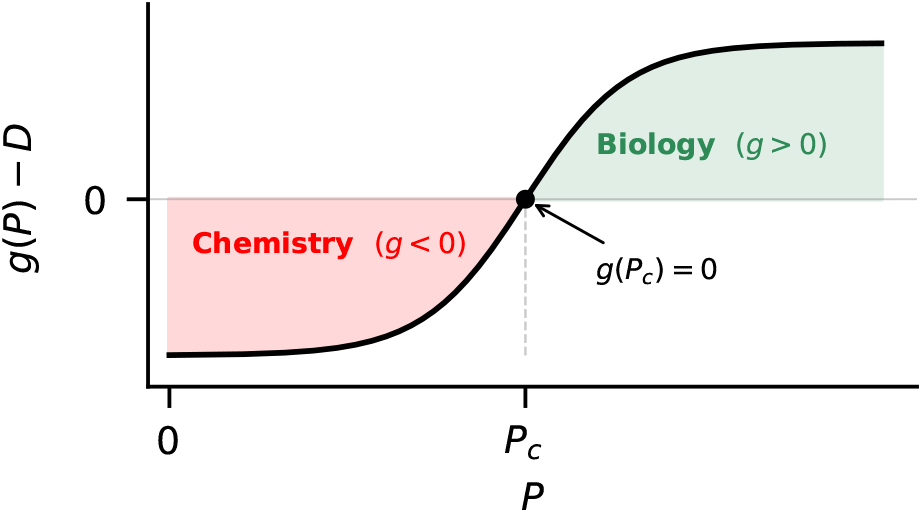
Cooperativity is required for the chemistry to biology transition: How the growth function (*g − D*)(*P*) depends on population *P* , which acts here as a reaction coordinate from chemistry to biology. (Left, prebiotic chemistry,) *g − D <* 0, degradation dominates over growth. (Right, biology,) *g − D >* 0, growth dominates over death. The transition requires cooperativity whereby a mechanism changes how the birth-death dynamics depends on population *P* .

### Proteins persist through folding and catalysis

First consider the protein component. What dynamical mechanism could explain the cooperativity/snowballing (i.e. the *g*_2_ term) for producing elongating protein chains from amino acids. We adopt here the *Foldcat mechanism (7, 9, 10)*. We take as given that amino acids could have been on the early earth (38–40), and that they could have undergone covalent linkages into short random peptides by prebiotic catalysts (41–43). While typical polymers have exponentially diminishing populations with chain length, the Foldcat hypothesis shows in computer modeling that heteropolymers, such as proteins, composed of hydrophobic (H) and polar (P) monomers, could elongate by two forms of cooperativity. Even random short HP chains will collapse into compact structures in water, reducing the degradation rates of their cores. And at a primitive level, such peptides could catalyze reactions such as chain elongation; see (Fig. 5), contributing to the *g*_2_ cooperativity term needed for advancing towards longer-chain biology (8). Importantly, it defines an early form of fitness, applicable to molecules in a prebiotic world; namely their relative persistence in the environment. If an ecumenical mechanism produces different monomer sequences, those molecules will have differential abilities to fold and catalyze, and not degrade, and thus persist longer.

**Figure 5:**
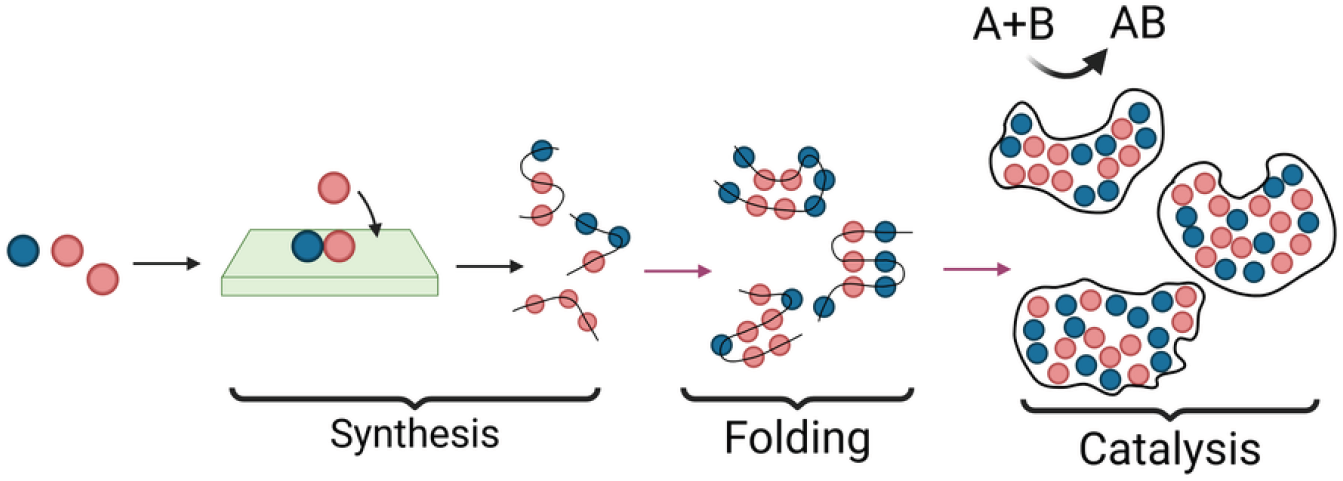
Foldcat mechanism: 1) Random HP chains are synthesized on a “founding rock”. 2) Short HP chains fold into stable structures due to hydrophobic collapse. 3) A subset of those folded structures serve to catalyze reactions by virtue of localized collections of sidechains.

But a proteins-only world is not sufficient to escape to biology. The Foldcat mechanism alone never leads to lineages of multi-molecule complexes. Also, the longer the protein chains become, the more difficult it is to further extend that chain because their folding buries their terminal ends, preventing further extension. At this point, it is important to distinguish what happens to a given HP sequence *i*, and its population *P*_*i*_, versus the population *P* of all sequences in the sequence space, such as in (2). Any single sequence will die out at death rate *D* by degradation and dilution (see SI), with dynamics

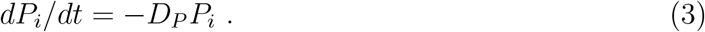

The mean lifetime of any specific sequence is proportional to 1*/D*_*P*_ , while the total population is governed by (2) and is able to exhibit cooperativity toward biology. Now consider how partnering of HP foldamer chains with RDNAs can give a huge advantage of fitness as persistence.

### RDNA gives fidelities, but lacks selection

A mechanism of RDNA would entail replication. As in the case with proteins above, we label sequences by the index *i*. Sequences have a birth and death rate, and its birth is dependent on having a template. However, because RDNA is ecumenical, no one sequence has any advantage over any other. For an RDNA-only world, each RDNA sequence has birth-death dynamics (see SI for details):

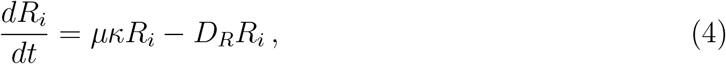

where *κ* is the replication speed which can be related to the doubling time of an RDNA population, having units of 1/time. And *µ* is the probability that a sequence makes an exact copy of itself (the replication fidelity), a dimensionless quantity representing the accuracy of correctly attaching a correct Watson-Crick pair versus an incorrect pair. In (4), what matters is only the product *µ × κ*, but below we need these two quantities separately. Finally, we note that in accord with our assumption above that the replication machinery itself must be agnostic to what sequence it is replicating, *κ* does not have an index *i* as we are assuming that heritable information is replicated, agnostic to the particular sequence that is being replicated.

In this model, RDNA is capable of self-copying, but has no mechanism for fitness/selection, because copying is assumed to be ecumenical, the same for all sequences. In addition, an RNA-world that contains only individual molecules lives only on a molecular timescale, since molecules degrade with prebiotic rates (44, 45).

### Central dogma replicator model

Our model below is based on the reasoning above. We assume a separation of actions whereby peptides fold and function as in the Foldcat model and they aid the replication of RDNA; see Fig. 6. We assume a system that can range from complete disorder to order. In the ordered state, RDNA guides protein synthesis; some such actions have been observed experimentally (46–49). Any such model requires some form of cooperativity, wherein the larger the protein population, the better the RDNA replication. Our model does not explicitly address the microscopic basis of this cooperativity, but various potential sources could be considered: activation of substrate; (50); positively charged proteins can assist the polymerization of negatively charged RDNA (30); of the electrostatic forces that tend to localize and stabilize RDNA-Protein biomolecular condensates (51). The model does not require unique folds. Biocondensates often have intrinsically disordered proteins that can catalyze multiple reactions (52–55). We don’t seek a more granular mechanism here, in order to avoid speculating and additional parameters that would obscure the main points here.

**Figure 6:**
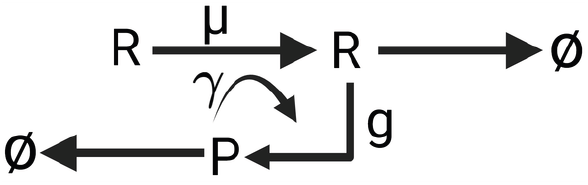
Protein-RDNA CD model: *µ* is a heritability order parameter for how well RDNA can copy itself. *γ* is a functionality/catalyst order parameter for how well proteins assist heritable copying. *g* is the rate at which RDNA molecules produce protein.

Assuming the two polymers have a CD relationship gives the following dynamics:

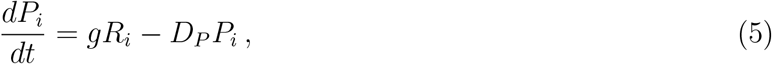

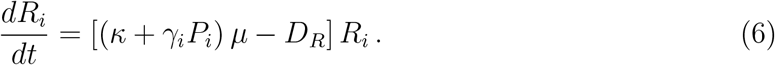

where *P*_*i*_ and *R*_*i*_ are the protein and RDNA populations respectively and the index *i* refers to individual sequences of RDNA and proteins. The coupling parameters *g* and *D*_*P*_ represent the rate of protein synthesis and degradation, and *γ* represents the protein fitness for the job of replicating any RDNA on a template.

Here are some properties of the model. When *g* = 0, proteins are not encoded by the RDNA, the protein population decays to zero. That situation reduces to an RDNA-only world. It does not exhibit cooperativity, all the RDNA sequences are equivalent, and they have no differential fitness. Instead, *γ* = 0 describes a situation where protein does not aid RDNA replication. This too has no cooperativity or selection. Only when *g* and *γ* are both non-zero does the model describe the organization of proteins and RDNA in an interacting assembly. This unit of evolution now has cooperativity since *γ*_*i*_, which depends on the sequence. And specific protein sequences are now able to control the growth of the assembly by indirectly controlling the RDNA population *R*_*i*_. In contrast, in a protein-only world, each sequence contributes to an overall average fitness 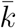, and then the label *i* is best regarded as the label of a specific assembly of proteins and RDNA.

A key conclusion is that this mechanism, (5) and (6), has the type of cooperativity needed according to (2) to allow the escape from chemistry to biology. Focusing on a specific sequence *i* we drop the index *i* from all quantities and define 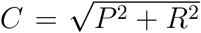, an effective quantity relevant to whether the assembly grows or decays. In the SI, we performed standard stability analysis and found that a transition from growth to decay appears in our system as a saddle node fixed point. Near the transition the growth/decay dynamics are given by:

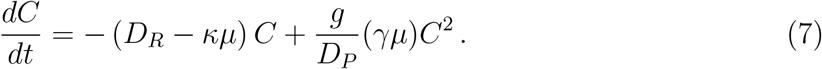

(7) has the cooperativity form of (2). It corresponds to growth rate *g*_1_ = *κµ*, degradation rate *D*_*R*_ and cooperativity parameter *g*_2_ = (*gγµ*)*/D*_*P*_. The linear growth term is entirely controlled by the RDNA replication parameters, consistent with the Central Dogma principle. Proteins only affect this transition via the parameter *γ*. Note that the cooperativity parameter is linear in the product *γµ* if we consider only the lowest-order coupling in a 2PW.

Now, we describe how (7) predicts a chemistry-to-biology transition. The protein fitness and RDNA fidelity parameters are captured through *γ* (proteins assisting RDNA copying) and *µ* (RDNA copying fidelity), respectively. Therefore, to address the Eigen paradox, we determine the relation between the critical value of these two order parameters required for escape. This is slightly different from the usual Eigen threshold (15, 16), which describes the stability of information in an error-prone replication system. Rather, we focus here on finding the threshold of fitness and fidelity needed for our model protein-RDNA assembly to transition toward biology.

We modeled the critical catalytic strength *γ*_*c*_ and fidelity *µ*_*c*_ to obtain the transition barrier of survival probability using the Gillespie algorithm and the deterministic boundary is obtained by solving (5) and (6) numerically, using the following parameters: *D*_*P*_ = 1, *D*_*R*_ = 0.5, *κ* = 0.05, *g* = 0.006, and (*P*_0_, *R*_0_) = (10, 10). The red region is where proteins and RDNA are not sufficient to escape chemistry; the green region is where the polymers are sufficient to escape. The blue curve on the right in Fig. 7 separating the red from green regions is the tipping line from chemistry to biology. The tipping line curve resembles a Pareto front where low fidelity RDNA can be compensated for by high fitness proteins or vice versa.

**Figure 7:**
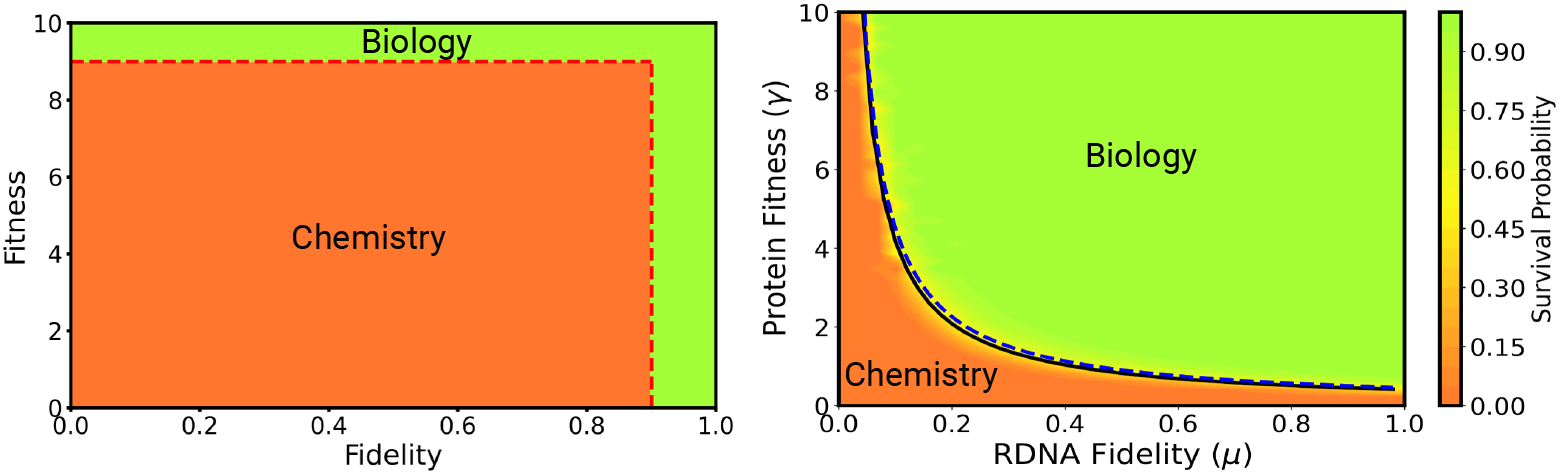
Computed Pareto Front for escaping from chemistry to biology, for the two evolutionary parameters *µ* and *γ*. (Left, Eigen box assumption:) both proteins and RDNA must be good independently to escape chemistry. The horizontal red line shows the necessary fitness in a protein-only world with poor fidelity *µ* = 0.111. The red vertical line is the necessary RDNA fidelity in an RDNA-only world with poor fitness *γ* = 1.11. (Right, computed phase diagram from the model:) (Red) Both values are small, growth dies out. (Green) The combination of values is large enough that escape to biology is autocatalytic. (Black solid line) The deterministic boundary obtained by numerically solving the system. (Blue dashed line) is our analytical approximation of the Pareto Front boundary.

The shape of this Pareto front can be obtained analytically by an informal argument. The critical point is determined by the parameters *g*_1_ and *g*_2_ in (2). Since *g*_1_ doesn’t depend on *γ* or *µ*, the relation between *γ*_*c*_ and *µ*_*c*_ can be obtained by essentially fixing the critical *g*_2_ ∝ *γµ*. Therefore, we obtain the functional form of the Pareto front *γ*_*c*_ ∝ 1*/µ*_*c*_. A more rigorous calculation is given in the SI which also allows the analytical determination of the Pareto front as follows:

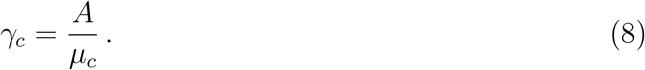

where *A* is a prefactor that depends on the remaining parameters and the initial concentrations *P*_0_ and *R*_0_. The exact form of *A* is given in the SI. The analytical computation of the Pareto front is valid in a prebiotic approximation, wherein protein synthesis is slower than protein degradation (*D* ≫ *g*). However, we find excellent agreement between the analytical solution and the numerical solution as shown in Fig. 7. In general, we expect the inverse relation between *γ*_*c*_ and *µ*_*c*_ to still hold even if this approximation is not valid, with only a change in the prefactor.

This model is simplified. It is only intended to capture the essence of the two sequence polymers, and assumes the CD relationship shown in Fig. 6. It is not a microscopic mechanism of their association. And while it does assume the dynamics of multi-molecule objects, it leaves out metabolites, cell-like encapsulation and small molecule processes that may have accompanied this 2-polymer association (56–59). We note, however that random RNA-protein associations are not rare events, as is becoming increasingly understood in studies of their biomolecular condensates (51, 60–70). In particular positively charged proteins naturally associate with RNAs, which are negatively charged in solution, across many different compositions, chain lengths and sequences (71).

Comparing the left and right panels in Fig. 7 shows the insight this model gives into the origins of a 2PW. The right figure shows what we call the *Eigen Box*. It supposes, as imagined in the Eigen Paradox, that good protein is needed to make RDNA, and that good RDNA is needed to make protein, and that the two polymers must achieve those conditions independently. For this figure, the definition of “good” is arbitrarily set to 90% of perfect. Users could choose other values, or could explore sequential mechanisms, one polymer coming before the other (72). Our model does not calculate the timescale for the emergence of this cooperativity. But this comparison demonstrates that a 2PW is more likely to emerge under the reasonable assumption that high fitness proteins are slower to emerge compared to low fidelity RDNA or vice versa. In short, the main point of comparison of the two panels of Fig. 7 is in showing that once RDNA and proteins become coupled – even if each one individually is poor at its job – the two polymers can cooperate in a sort of double-bootstrapping: poor-quality RNA can be rescued by good-quality protein or vice versa.

Fig. 8 shows an important conclusion from the model; namely that the joining together of RDNA with protein leads to far greater persistence – and thus far greater fitness – than is achievable by either individual polymer alone. This is because when RDNA and protein combines cooperatively to evolve with some degree of heritability – even with only moderate fidelity – those complexes now have the much longer lifetimes as lineages than the lifetimes that individual molecules alone would have. Fig. 8 shows the different levels of persistence with different model parameters. The curve with *γ* = 0, no protein assistance for RDNA replication, shows little ability to persist for any value of *µ*, the fidelity parameter. In contrast, the curve with *γ* = 2, representing some degree of coupling, reaches much longer persistence, for sufficiently large *µ*. In comparison, the molecular lifetime is not affected by the coupling, meaning that this fitness advantage is not due to the increase in the lifetime of any single RDNA molecule, but due to the cooperativity provided by proteins. These plots were generated with an initial population of 6 RDNA molecules and a maximum simulation time of *t* = 100. Therefore the plateau occurring after the dashed vertical line is an artifact of our finite simulation time.

**Figure 8:**
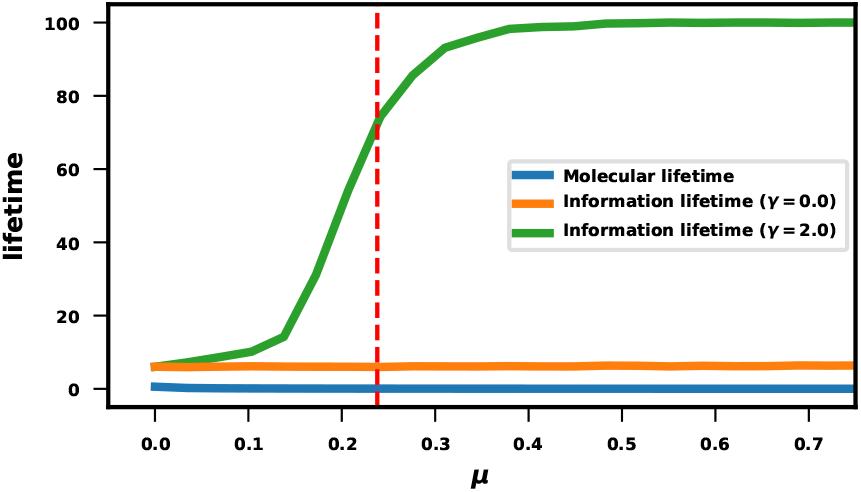
The persistence (fitness) is much greater for Protein-RDNA than for either alone. Plots showing the lifetime of the entire RDNA population (the information lifetime) and the average lifetime of a single RNA molecule in the system as a function of *µ* for *γ* = 0 (orange curve), and *γ* = 2.0 (green curve).

## 1 Summary

How might prebiotic chemistry have led RDNA and proteins into cooperative functional associations? We give here a model that assumes a separation of labor, proteins acting to assist RDNA replication fidelity, in a central dogma arrangement. The model has two evolutionary parameters, allowing us to study the emergence of order from disorder. The model has the type of second-order cooperativity needed to get from chemistry to biology. The two polymers leverage each other to grow to high joint fitness, where lineages now dominate over single molecules. The model predicts a double bootstrapping cooperativity: This 2-polymer world (RDNA, protein) had substantially better ability to persist than either single polymer world alone would have had.

## Code Availability

The code used to conduct this research can be found in a public github repository at: https://github.com/mswailem/RP_CD_model

## Acknowledgments

The authors are deeply grateful to Charles Kocher, Ying-Jen Yang, Lakshmanji Verma, Anthony Bogetti, Ron Zuckermann, Donghui Zhang, and Samuel Owoso for fruitful and thoughtful discussions regarding this work. We appreciate support from the Laufer Center for Physical and Quantitative Biology and the National Institute of General Medical Sciences of the National Institutes of Health under award number RM1GM135136.

