## Supplementary Information for "RNA and proteins joined up at the origins of life: Persistence is the point"

Mohamed Swailem
Ken A. Dill

July 9, 2026

#### The Central Dogma precludes Lamarckian evolution

If evolution were Larmarckian, rather than Darwinian, it would violate the Central Dogma (as given by Fig. 2 in the main text). In the Lamarckian evolution model, interaction and experience of the organism with its environment are able to lead to changes in the genome ("Giraffes grow taller so that they can reach the leaves as food in high trees."). In principle, the assembly line improvement mechanism in the text could be done either in a Larmarckian or Darwinian way. The Central Dogma says that all information flows from the blueprint. So, when a mutation is proposed in Darwinian evolution, it is proposed directly on the blueprint $N$, namely in the DNA, before competition determines if the proposal would be a successful change. So, whichever is the winner cell – either the wildtype or mutant – already carries that information into the next generation. The winner has blueprint $N^x$. In contrast, for Lamarckian change, instead of mutating in N, you would mutate in P directly. Then, after discovering which P has the best fitness, you would modify N accordingly after the fact. Here's the Lamarckian idea in the blueprint metaphor. Imagine that at the end of the competition process, the winner is learned to be $P^x$. Then the Lamarckian mechanism would require a *Reverse Dogma* step: READING the $P^x$ and WRITING it into $N^x$. Why couldn't that happen? As we have argued above, polymer physics doesn't allow it because proteins are largely not readable. Lamarckian evolution is incompatible with the Central Dogma, and it is because of the different physics of N and P polymers.

#### Sequence decomposition of single polymer worlds.

A protein-only polymer world settles into a stationary distribution of chain length given by a Flory-like distribution (1). Let's decompose the general peptide population $P$ into a population of sequences $P_i$, where $i$ is the sequence label. Thus we have $P = \sum_{i=1}^{N} P_i$, with $N$ being the size of the sequence space. Given that the foldcat mechanism enables the formation of long peptide chains capable of folding and function as self-catalysts, we can assume a large sequence space $N >> 1$. The dynamics of the population decomposition is modeled using Eigen's model of molecular evolution (2):

$$\frac{dP_i}{dt} = \sum_{j=1}^{N} Q_{ij} k_j P_j - D_P P_i \,, \tag{S1}$$

where $Q_{ij}$ is the mutation matrix describing the formation of the sequence $i$ from the sequence $j$, and $k_j$ is the fitness contribution of sequence $j$ to the overall foldcat growth function $g(P)$. Since a replication event forms another peptide sequence, the mutation matrix needs to obey the following identity: $\sum_i^N Q_{ij} = 1$. It can be shown, using the mutation matrix identity, that the total population evolves over time according to the following equation:

$$\frac{dP}{dt} = \bar{k}(P)P - D_P P \,, \tag{S2}$$

where $\bar{k} \equiv \sum_i^N k_i P_i / P$ is the average fitness of the population. Comparing this with Eq. (2) in the main text, we make the identification $g(P) = \bar{k}(\mathrm{P})$. Note that the fitness of each individual sequence need not be a constant, but can depend on the total population and the sequence distribution. Peptides cannot act as templates for the synthesis of other peptides as dictated by the Central Dogma. Therefore, the mutation matrix is a result of random peptide chain synthesis: $Q_{ij} = 1/N$, which upon performing a large $N$ approximation gives:

$$dP_i/dt = -D_P P_i \ . \tag{S3}$$

This equation shows that in a protein-only world, cooperativity can exist on a population level, but each individual sequence or a finite network of Eigen hypercyclic sequences (relative to the large $N$ approximation) is always stuck in the degradation dominated chemistry phase.

The RDNA dynamics are also modeled using Eq. (S1) under the assumption that the fitness $\kappa$ is dominated by a universal replication mechanism, and therefore does not carry any sequence dependence at lowest order. This assumption follows from the physical arguments for the Central Dogma as discussed in the main text. The sequence decomposition of an RDNA population is then given by:

$$\frac{dR_i}{dt} = \sum_{j=1}^{N} \tilde{Q}_{ij} \kappa R_j - D_R R_i \,, \tag{S4}$$

where $\kappa$ is the fitness of the universal replication mechanism and $D_R$ is the degradation rate. The RDNA-world mutation matrix $\tilde{Q}_{ij}$ now depends on Watson-Crick pairing dynamics (3). For our level of modeling, it suffices to parametrize it through the probability $\mu$ of a sequence successfully making an exact copy (The replication fidelity), and we assume that once a mutation occurs, the resultant mutant is selected uniformly from the sequence space. A more accurate level of modeling would take the Hamming distance between sequences into account (4), however, this complicates the mathematical modeling without changing our conclusion. After these considerations, the RDNA mutation matrix is given by:

$$\tilde{Q}_{ij} = \begin{cases} \mu & i = j \\ (1-\mu)/N & i \neq j \end{cases} \tag{S5}$$

Using this form and assuming $N \gg 1$, then each sequence obeys the equation:

$$\frac{dR_i}{dt} = \mu\kappa R_i - D_R R_i \, . \tag{S6}$$

This is the equation we use in the main text to investigate an RDNA-only world.

### Stability analysis and decomposition of the Central Dogma dynamics

In this section, we perform the standard stability analysis of the dynamics in Eq. (5) and Eq. (6) in the main text and reduce it to a one-dimensional system that can undergo an invasion analysis that is performed in (1).

First, we identify the fixed points (FP) of the system:

- The extinction FP: $(0,0)$, which is always stable iff $D_R - \kappa\mu > 0$.
- The saddle point: $(P^* = [D_R - \kappa\mu]/[\gamma\mu]; \;\; R^* = (D_P P^*/g)$, which is always a saddle point and only exists in the positive region of phase space iff $D_R - \kappa\mu > 0$.
- Escape to infinity: $(P \to \inf, R \to \inf)$, which is always stable.

The unstable manifold can be approximated near the saddle point by finding the eigenvalues and the eigenvectors of the Jacobian matrix at that point:

$$\epsilon_+ = \frac{1}{2}\left(-D_P + \sqrt{D_P^2 + 4D_P\big(D_R - \kappa\mu\big)}\right) \qquad \Longrightarrow \qquad \mathbf{v}_+ = \begin{pmatrix} g \\ \epsilon_+ + D_P \end{pmatrix} \, , \tag{S7}$$

$$\epsilon_- = \frac{1}{2}\left(-D_P - \sqrt{D_P^2 + 4D_P\big(D_R - \kappa\mu\big)}\right) \qquad \Longrightarrow \qquad \mathbf{v}_- = \begin{pmatrix} g \\ \epsilon_- + D_P \end{pmatrix} \, , \tag{S8}$$

where $\epsilon_+ > 0$ corresponds to the unstable eigenvector and $\epsilon_- < 0$ corresponds to the stable eigenvector. These eigenvectors provide linear approximations for the unstable manifold and the separatrix near the saddle FP. Near the transition point we approximate the unstable manifold by a linear curve passing through $(0,0)$ and the saddle FP for large $D_P$ values (which is the prebiotically relevant situation), and for small $P$ and $R$ concentrations around the FP. A transformation into polar coordinates $(P,R) \to (C = \sqrt{P^2 + R^2}, \tan\phi = R/P)$ naturally decomposes the dynamics into composition dynamics controlled by the polar angle

$\phi$ and growth/decay dynamics controlled by the "radial" coordinate $C$. We are interested in the behavior of the growth/decay dynamics, which is given by:

$$\frac{dC}{dt} = \frac{P}{C}\frac{dP}{dt} + \frac{R}{C}\frac{dR}{dt} \tag{S9}$$

Substituting this into Eq. (5) and Eq. (6):

$$\frac{dC}{dt} = \left(-D_P\cos^2(\phi) + g\cos(\phi)\sin(\phi) - (D_R - \kappa\mu)\sin^2(\phi)\right)C + \gamma\mu\cos(\phi)\sin(\phi)C^2\,. \tag{S10}$$

Therefore the dynamics of the progress coordinate $C$ consist of a linear growth term and a cooperativity $C^2$ term. So far, this equation has been exact. The issue is that in general $\phi(t)$ changes over time, and so the dynamics of $C$ are not closed, as they couple to the dynamics of $\phi$. However, as we are interested in the behavior of the system near the transition point, we examine the dynamics of $C$ near the saddle node FP by assuming that $\phi$ is constant and given by the relation $\tan(\phi) = R^*/P^* = D_P/g$. From this relation we can find:

$$\sin(\phi) = \frac{D_P}{\sqrt{D_P^2 + g^2}} \approx 1\,, \tag{S11}$$

$$\cos(\phi) = \frac{g}{\sqrt{D_P^2 + g^2}} \approx \frac{g}{D_P}\,. \tag{S12}$$

Where we have used the approximation $D_P >> g$, which is appropriate in a prebiotic context, where protein synthesis is slow compared to protein degradation. Substituting this into (S10), we obtain:

$$\frac{dC}{dt} = -\left(D_R - \kappa\mu\right)C + \frac{g}{D_P}(\gamma\mu)C^2\,. \tag{S13}$$

This equation has the same form as Eq. (2) with growth rate $g_1 = \kappa\mu$, degradation rate $D_R$ and cooperativity parameter $g_2 = g\gamma\mu/D_P$.

### Analytical determination of the Pareto front

In order to analytically determine the Pareto front shown in Fig 7 in the main text, we first write Eq. (5) and Eq. (6) in non-dimensional form using the transformations:

$$P \to x \equiv \frac{P}{P^*}\,, \tag{S14}$$

$$R \to y \equiv \frac{R}{R^*}\,, \tag{S15}$$

$$t \to \tau \equiv D_P t\,. \tag{S16}$$

This leads to the following non-dimensional equations:

$$\frac{dx}{d\tau} = y - x\,, \tag{S17}$$

$$\frac{dy}{d\tau} = r(x-1)y\,. \tag{S18}$$

where $r \equiv (D_R - \kappa\mu)/D_P$ is the ratio between the loss of $R$ to the loss of $P$. The Pareto front can be determined if an analytical expression for the separatrix is found.

The separatrix curve is given by $y_s(x)$ which is the curve in the $(x, y)$ phase space (or equivalently the $(P, R)$ phase space) that separates solutions that flows to the extinction FP from solutions that escape to infinity. The separatrix curve satisfies the following equation:

$$\frac{dy}{dx} = r\frac{(x-1)y}{y-x}\,, \tag{S19}$$

with boundary condition $y_s(x = 1) = 1$. Initial conditions that lie above the separatrix $y(t = 0) > y_s(x(t = 0))$ will escape, while solution below it $y(t = 0) < y_s(x(t = 0))$ will decay into extinction. Therefore, the Pareto front is determined by the condition $y(t = 0) = y_s(x(t = 0))$. A closed form solution for the above equation is unobtainable. However, due to the transformation:

$$y = \frac{R}{R^*} = \frac{g}{D_P}\frac{R}{P^*} = \frac{g}{D_P}\frac{R}{P}x\,. \tag{S20}$$

If we assume that the initial concentration of $P$ and $R$ are of the same order of magnitude, and in a prebiotic environment $g/D_P << 1$, then $y/x \sim (g/D_P)O(1) << 1$. In this approximation regime we can find a closed form solution to Eq. (S19) as follows:

$$\frac{dy}{dx} \simeq r\frac{1-x}{x}y\,, \tag{S21}$$

$$y_s(x) = x^r e^{r(1-x)}\,. \tag{S22}$$

Where the boundary condition $y_s(x = 1) = 1$ was used. Using this solution alongside the Pareto front equation $y(t = 0) = y_s(x(t = 0))$ and transforming back to the $P$ and $R$ coordinates, we obtain (after algebraic manipulation):

$$(\gamma\mu)^{1-r}\, e^{P_0\mu\gamma/D_P} = \frac{D_P^2}{g}\left(\frac{P_0}{D_P r}\right)^r \frac{r}{R_0}e^r\,. \tag{S23}$$

Where $P_0$ and $R_0$ are the initial concentrations of $P$ and $R$, respectively. Since $r$ depends on $\mu$, this represents a complicated functional dependence of $\gamma(\mu)$, in order to simplify this expression we take the prebiotically relevant approximation of large degradation rates $r = (D_R - \kappa\mu)/D_P \sim D_R/D_P$, and therefore does not depend on $\mu$. Then $\gamma$ can be obtained using the Lambert W function and we find:

$$\gamma(\mu) = \frac{A}{\mu}\,, \tag{S24}$$

where:

$$A \equiv \frac{D_P - D_R}{P_0} W\left[\frac{D_R}{D_P - D_R}e^{\frac{D_R}{D_P - D_R}}\left(\frac{D_P}{g}\frac{P_0}{R_0}\right)^{\frac{D_P}{D_P - D_R}}\right]\,. \tag{S25}$$

This approximation of the Pareto Front works when the initial conditions are such that $y << x$ (or equivalently $(gR_0)/(D_P P_0) << 1$). However, an analytical approximation of the

Pareto Front in the regime $x >> y$ (or equivalently $(gR_0)/(D_P P_0) >> 1$) is also permissible by determining the separatrix in this regime:

$$\left.\frac{dy_s}{dx}\right|_{x>>y} = -r y_s \,, \tag{S26}$$

which can be easily solved with the boundary condition $y_s(1) = 1$, leading to:

$$y_s(x) = 1 + r - rx \,. \tag{S27}$$

Applying the same condition as with the $y << x$ case, and transforming back into the $P$ and $R$ coordinates, the Pareto Front in this regime is determined as:

$$\gamma(\mu) = \frac{1}{\mu}\frac{D_R(D_P + D_R)}{gR_0 + D_R P_0} \,, \tag{S28}$$

where we have used the large degradation approximation $r \sim D_R/D_P$ and $P^* \sim D_R/(\gamma\mu)$.
